# Phage integrons are vaults of privatized counterdefense genes

**DOI:** 10.64898/2026.11.24.768926

**Authors:** Ryuichi Ono, Brady F. Cress

**Author notes:** Corresponding author. (B.F.C.).

## Abstract

Integrons facilitate the adaptation of microbes by capturing beneficial gene cassettes, including antiphage defense systems, but their role in phage adaptation remains unclear. Here, we identify reversed integron cassette organization in phages, with 96% of reverse-oriented cassettes lacking detectable homologs in surveyed bacterial and plasmid integrons. Phage cassettes are enriched in counterdefenses, including viral sponge-like proteins. Despite these differences, representative phage and bacterial integron integrases recombine cassette attachment sites from either source; however, exchange between oppositely organized phage and bacterial integron arrays misorients incoming gene cassettes relative to the recipient integron’s promoter, posing an expression barrier. These findings establish integrons as a shared engine of the phage-host arms race, capturing defenses in bacteria and counterdefenses in phages, while cassette orientation buffers exchange between the two adaptive repertoires.

## Introduction

The arms race between prokaryotes and their mobile genetic elements (MGEs) is one of the most ancient and strongest driving forces of their evolution (*1*). This evolutionary pressure is clearly illustrated by bacterial systems that assemble and renew their defensive repertoire; one well-characterized example is CRISPR, which adapts to changing targets by capturing fragments of invading DNA into its array. Recent studies have shown integrons to be another example of the adaptive microbial defensive armory. Integrons are genetic machinery that accumulate beneficial genes, called cassettes. The integron integrase (IntI) facilitates recombination between *att* sites, leading either to cassette insertion (*attI*-*attC*) or excision (*attC*-*attC*), in an *attC* hairpin strand dependent manner (Fig. 1A). Together, insertion and excision enable hosts to take up beneficial genes and, in combination (excision followed by re-insertion at *attI*), to shuffle cassettes, bringing a cassette to the first position of the array next to the cassette promoter (P_c_). This fine-tunes expression levels and lowers the metabolic burden by expressing costly genes only when necessary (*2*). It is therefore reasonable that integrons, which keep genes in a regulated reserve, can stockpile defense systems that confer large fitness gains under appropriate control but are otherwise costly or even deleterious (*3*). Indeed, recent discoveries of diverse defense systems in integron cassettes support this nature as a defense system biobank (*4*–*7*).

**Fig. 1.**
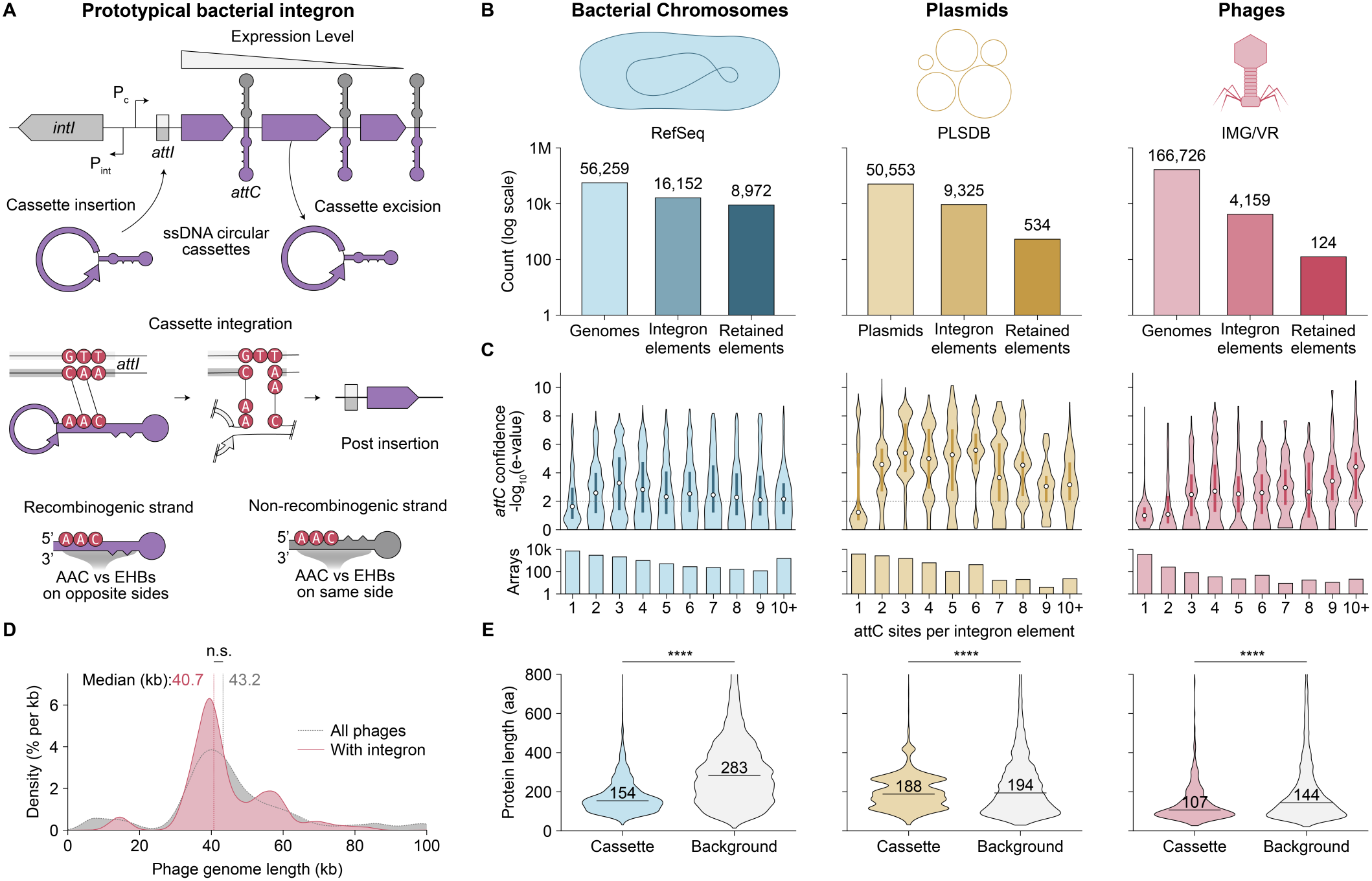
Identification of phage-encoded integrons in large-scale datasets. **(A)** Prototypical bacterial integron: integrase gene (*intI*) with promoter (P_int_), cassette promoter (P_c_), and cassettes (arrows) followed by *attC* hairpins. IntI inserts circular single-stranded cassettes at *attI* and excises cassettes. The AAC triplet and extrahelical bases (EHBs) from the recombinogenic strand of *attC* lie on opposite sides of the hairpin. **(B)** Sequences analyzed, integron elements with at least one *attC*, and elements retained after the *attC* threshold and deduplication, in bacterial chromosomes (CI), plasmids (MI), and phages. **(C)** Top, *attC* detection confidence (−log10 E-value) by the number of *attC* sites per element (white dots, medians; thick bars, interquartile ranges; dotted line, E-value of 0.01). Bottom, number of elements (arrays; log scale). **(D)** Genome length of all IMG/VR phages (n = 166,726) and of the 117 phages with integron arrays of at least three *attC* sites. Dotted lines, medians; n.s., not significant (Mann-Whitney U test); genomes over 100 kb are not shown. **(E)** Length of cassette (left) and background (right) proteins. Lines and numbers, medians; ****, *P* < 10^−4^ (Mann-Whitney U test); proteins longer than 800 aa are not shown.

In contrast to our increasing understanding of defensive armory formation, the other side of the arms race—the formation of the phage counterdefense armory—is largely uncharacterized. The prototypical role of integrons as a defensive biobank and a recent report of phage-encoded integrons (*8*) lead to an exciting hypothesis: phage-encoded integrons may serve as a biobank of counterdefense genes. However, the number of phage-encoded integrons found so far is very limited (20 elements, only 3 of which encode both integrase and *attC*). Moreover, no hallmark features exist that distinguish these elements from other integron subtypes, leaving the existence and function of phage-specific integron subtypes an unanswered question.

Here we show that phage-encoded integrons are a distinct integron subtype with a unique architecture and role as a counterdefense biobank. Phage-encoded integrons streamline phage genome evolution without expanding the genome size by capturing miniature cassettes. Their cassettes mostly lie in the reverse orientation compared to non-phage integrons, and their cassette pool is largely distinct and characterized by an enrichment of counterdefense-related functions, including anti-restriction. Further structure-guided mining and experimental screens identified functional anti-signaling molecule sponge proteins, highlighting the role of phage integrons as a counterdefense biobank. Despite being associated with reverse cassettes, a representative integron integrase of the phage-specific clade has the same strand specificity as the class 1 integron integrase. Because of this shared specificity, reverse and forward cassettes insert in opposite orientations, therefore exchange between phage and non-phage integrons misorients incoming cassettes relative to the recipient integron’s promoter, posing an expression barrier that would privatize the cassette pool. These cassettes, which we named Cassettes In Phage Integrons Held in Exclusive Reserve (CIPHERs), suggest a role for phage-encoded integrons as a privatized biobank of counterdefense genes. Our work establishes phage-encoded integrons as a distinct subtype of mobile integron with a likely origin independent from widespread class1 mobile integrons, and as a system to streamline the development and renewal of counterdefense armory in phage genomes.

## Results

### Discovery of phage-encoded integrons from large-scale datasets

Through stringent searching and data curation of 166,726 high-quality phage genomes from IMG/VR, we successfully collected 124 high-confidence, non-redundant phage-encoded integron elements (Fig. 1B; see Methods). As a reference for known integrons, we analyzed bacterial chromosomes from RefSeq (chromosomal integrons, CI) and plasmids from PLSDB (mobile integrons, MI) in the same way. The number of cassettes (counted by the number of *attC*s) was consistent with previous reports for bacteria and plasmids (Fig. 1C): Most MI carried 1-6 cassettes (*2*), whereas arrays from CI tended to be longer, up to 331 (found in *Vibrio navarrensis*) (*9*). In phage-encoded integrons, 87.8% of elements contained a single *attC* and 5.9% contained three or more (Fig. 1C). Since cassette arrays with fewer *attC* sites may have resulted from false-positive *attC* detections, we set a threshold for the minimum *attC* count per integron element by examining the distribution of *attC* detection confidence (E-value) as a function of the number of *attC* sites in the array (Fig. 1C). In CI and MI, the median e-value of *attC* sites from elements with two or more *attC* sites was below 0.01. In phages, however, the median e-value of *attC* sites from elements with one or two *attC* sites was above 0.01, indicating either a substantially higher false-positive rate or the presence of divergent phage *attC*s. To guard against false positives, we considered only elements with at least two (CI and MI) or three (phage) *attC* sites in all subsequent analyses.

Gene accretion through integron capture seems at odds with the capsid-imposed limit on phage genome size, so we asked whether phages carrying integron arrays have larger genomes and how long their cassettes and arrays are. The 117 phage genomes carrying these 124 arrays had a median length close to that of all phage genomes in our dataset (Mann-Whitney U, *P* = 0.25; Fig. 1D). Cassette proteins were shorter than the non-cassette background proteins in all three datasets (154 vs 283 aa in CI, 188 vs 194 aa in MI and 107 vs 144 aa in phage; Mann-Whitney U, *P* < 10^−300^, 2.6 × 10^−6^ and 1.7 × 10^−30^, respectively; Fig. 1E), and phage-encoded integrons biased towards few numbers of cassettes relative to CI/MI, in line with the compressed cassette protein lengths reported previously (*6*). These results suggest that phages can accommodate gene capture by exploiting the small size of cassettes.

### Phage integrons carry reverse cassettes with a distinct gene repertoire

Phage-encoded integrons differ from known integrons in the architecture of their cassette arrays. Because *attC* acts as a single-stranded hairpin with only one recombinogenic strand (Fig. 1A), the start codon of an inserted cassette faces toward or away from *attI* depending on the orientation of its coding sequence (CDS) relative to that strand (Fig. 2B). We found that in most CI and MI (94.0% and 93.8% of cassettes, respectively), the recombinogenic strand of an *attC* and its adjacent cassette lie on the opposite strand (forward cassettes; Fig. 2A-C), whereas in phage-encoded integrons they mostly lie on the same strands (reverse cassettes; 70.4% of cassettes in phage-encoded integrons; Fig. 2A). Representative examples of a CI, an MI and a phage-encoded integron discovered here are shown in Fig. 2D-F. Toxin-antitoxin (TA) cassettes found in CI deserve special mention. They account for approximately 45% of the reverse cassettes in CI (Fig. S1) and are known to contribute to the maintenance of CI cassette arrays (*10*). Because TA cassettes carry a promoter independent of P_c_ (*11*), they can maintain transcription regardless of their orientation relative to P_c_ and are therefore thought to persist stably as reverse cassettes. In contrast, among the reverse cassettes of phage-encoded integrons, only one (a CcdB-family toxin) had a homolog of these TA cassettes. More generally, only 4% of the reverse cassettes had homologs in CI or MI (compared with 54.3% of the forward cassettes in phage-encoded integrons). These results suggest that reverse cassettes are an architecture characteristic of phage-encoded integrons, and that their cassette pool is largely distinct from CI/MI cassettes.

Together, these observations suggest a model in which phage-encoded integrons utilize reverse cassettes, CI use them only occasionally (as TA cassettes), and MI probably rarely use them. We supported this model using the e-values of *attC* predictions from the *attC* covariance model (Fig. S2; Methods). In CI and phage-encoded integrons, the *attC*s of both forward and reverse cassettes derived from high-confidence *attC* predictions, and those of reverse cassettes were, if anything, more confident. In MI, however, the *attC*s of cassettes predicted to be reverse derived from far lower-confidence predictions than those of forward cassettes, suggesting that many of them are false positives.

**Fig. 2.**
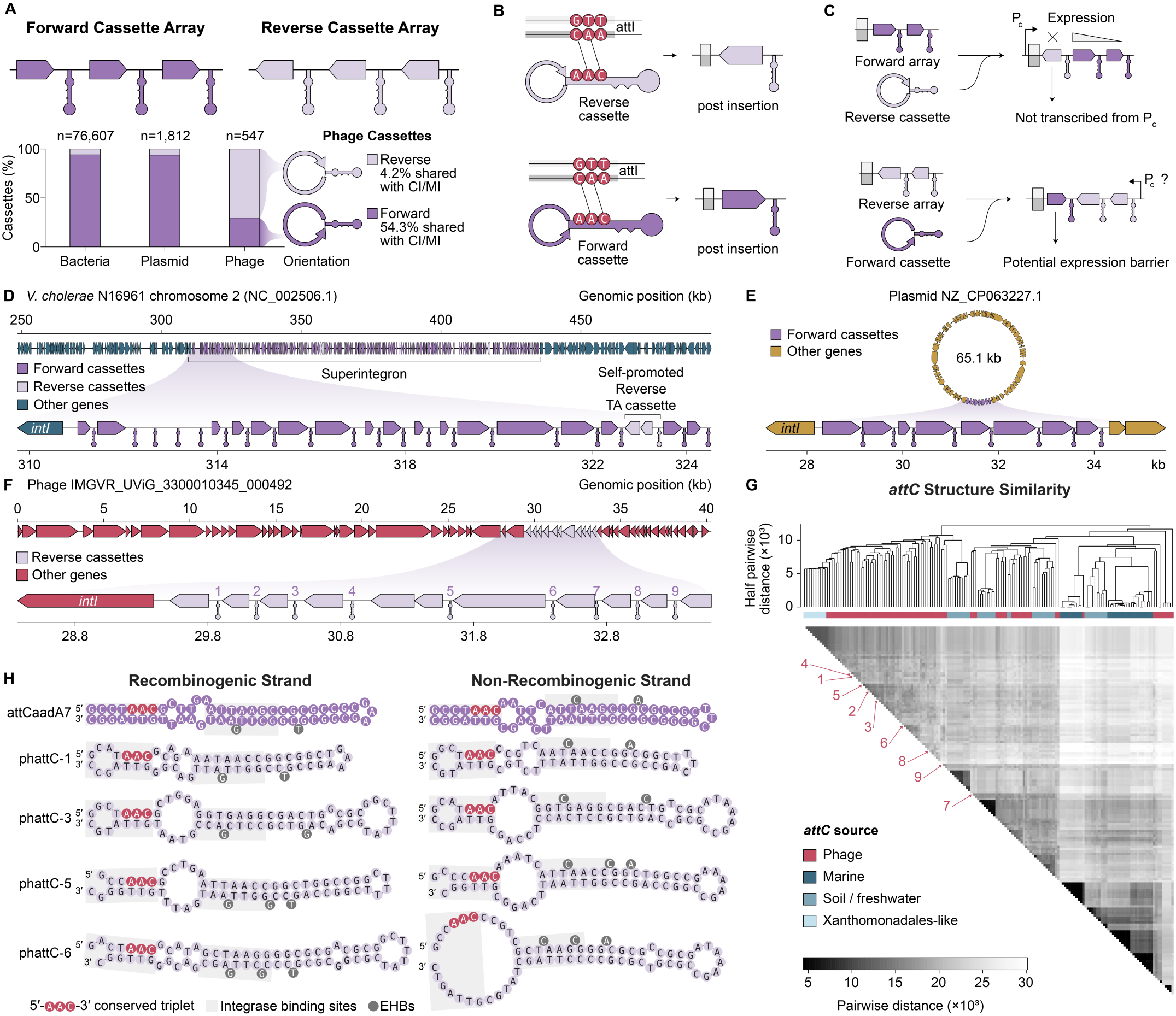
Phage-encoded integrons carry reverse cassettes with *attC*s of diverse origins. **(A)** Forward and reverse cassette arrays, defined by the strand of the CDS (arrows) relative to the recombinogenic strand of its *attC*. Bars, forward and reverse cassettes paired with an *attC* in bacterial chromosomes (CI), plasmids (MI), and phages. **(B)** Forward and reverse cassettes insert at *attI* in opposite orientations. **(C)** A cassette entering the other array type faces an expression barrier. **(D)** Superintegron of *Vibrio cholerae* N16961 chromosome 2. **(E)** Class 1 integron of the *Enterobacter hormaechei* plasmid NZ_CP063227.1. **(F)** Phage IMGVR_UViG_3300010345_000492 (*attC* sites numbered). In **(D)** to **(F)**, arrows are CDSs and hairpins are the recombinogenic strand of *attC*s; lower, integron region. **(G)** Structural clustering of 82 phage *attC*s (reverse cassettes; one per viral operational taxonomic unit, vOTU, plus all nine in D) and 80 reference *attC*s (Ghaly et al. 2021). Numbers, the *attC*s in **(F). (H)** Predicted structures of the recombinogenic (left) and non-recombinogenic (right) strands of four phage *attC*s (ph*attC-1, -3, -5, and -6*; *attC* 1, 3, 5, and 6 in F) and *attC-aadA7*.

We then asked where the *attC*s of phage-encoded integrons originated from. We structurally aligned the *attC*s of phage-encoded integrons (ones from reverse cassettes) with taxon-specific representative *attC*s (*12*) (Fig. 2G). The phage-encoded *attC*s were most often closest to the XL clade, which that study defined from Xanthomonadales *attC*s and whose environmental or taxonomic basis it left unresolved (49 of 82 *attC*s); 23 *attC*s were closest to the soil/freshwater clade and 10 to the marine clade. A single phage-encoded integron array could carry *attC*s from multiple sources (Fig. 2F; of nine *attC*s, seven were closest to XL and two to soil/freshwater), and the structures of these *attC*s retained the features of classical *attC*s, including conserved AAC triplet and Extrahelical bases (EHBs) (Fig. 2H).

### Activity and independent origin of a phage-encoded integron integrase

Since phage-encoded *attC*s derive from diverse sources, we hypothesized that their *intI*s also follow the same trend. Surprisingly, the phylogenetic tree of *intI*s containing the 10 *intI*s found in our dataset and diverse reference *intI*s from a previous study showed otherwise (*13*) (Fig. 3A; Methods). Six of the 10 *intI*s were accompanied by reverse arrays and four by forward arrays. Five of the six *intI*s with reverse arrays formed a single clade (SH-like approximate likelihood ratio test, SH-aLRT = 100) with an *intI* from a prophage in Betaproteobacteria, and four of these five phages were predicted by iPHoP (*14*) to infect Betaproteobacteria, namely Rhodocyclaceae. This clade was a sister clade of the one consisting of *intI*s from Betaproteobacteria, and the clade containing the clinically important class 1 and class 3 integron *intI*s was in turn sister to both. Mobile class 1 integrons are thought to have arisen through domestication of a Betaproteobacterial chromosomal integron by Tn402 (*15*). Phage-encoded integrons with reverse arrays thus appear to have an origin independent of such transposon-mediated capture events by plasmids: their *intI*s fall outside the clade of class 1 and class 3 *intI*s, and both the phylogeny and the predicted hosts suggest Betaproteobacterial chromosomal integrons as their direct source.

**Fig. 3.**
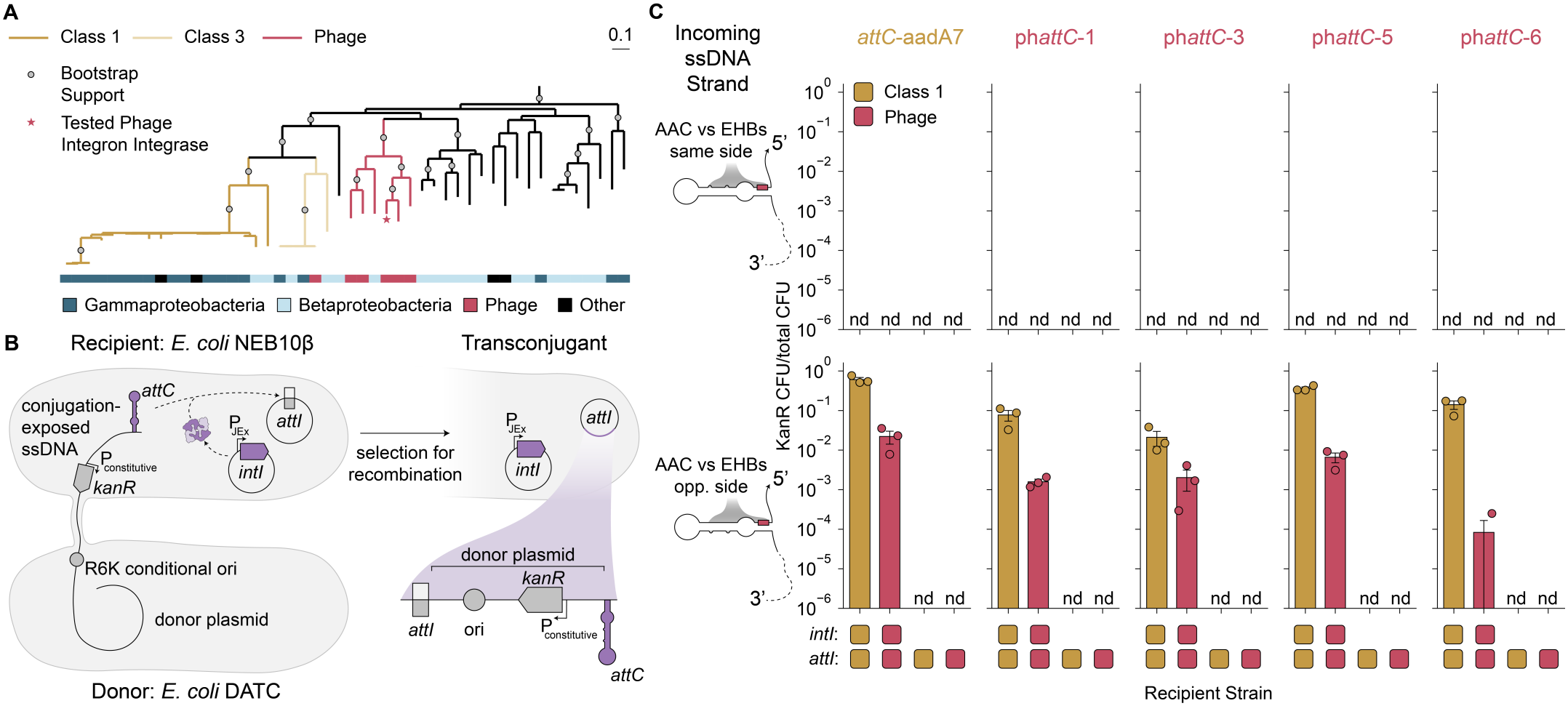
Phylogenetic position and activity of the phage-encoded integron integrase. **(A)** Maximum-likelihood phylogeny of integron integrases (187-sequence tree pruned for display; XerC/XerD outgroup). Clades of class 1, class 3, and reverse-array phage integrases are colored as indicated; the star marks the phage *intI* tested in **(C)**; circles mark SH-aLRT ≥ 95; the bar shows the taxonomy of the source organism. **(B)** Suicide conjugation assay. A donor *E. coli* (DATC) delivers a plasmid with an R6K origin (replication requires the π protein), an *attC* site, and KanR as single-stranded DNA to recipient *E. coli* NEB10β carrying *attI* and inducible *intI* on separate plasmids; only *attI*-*attC* recombination yields kanamycin-resistant transconjugants. **(C)** Recombination frequency (KanR CFU per CmR recipient CFU) for *attC-aadA7* and four phage *attC*s (*attC* 1, 3, 5, and 6 in Fig. 2F) delivered as the top strand (ts, upper; non-recombinogenic) or bottom strand (bs, lower; recombinogenic). The donor plasmids for each strand are listed in Table S1. Bars, left to right: class 1 *intI* with *attI1*, phage *intI* with the putative phage *attI*, and controls without *intI*. Mean ± SEM of three biological replicates (circles); nd, not detected.

We tested the activity of the phage-encoded *intI* with a suicide conjugation assay (Fig. 3B). All *attC*s whose structures are shown in Fig. 2H were tested as both the recombinogenic and the non-recombinogenic strand (Table S1). As positive controls, we used the class 1 *intI* (*intI1*) with its cognate *attI* site and *attC-aadA7* from previous studies (*16*). For the phage *intI*, we used the intergenic region (IGR) downstream of *intI* in the genome map shown in Fig. 2F as the putative *attI* site. For the recombinogenic strand of every *attC, intI1* showed recombination frequencies between 10^−2^ and 10°, and the phage *intI* showed lower but strand-specific and *intI*-dependent integration activity (Fig. 3C). These data show that the phage integron-specific *intI* and the putative *attI* site have cassette integration activity and that, although accompanied by reverse cassettes, this *intI* preserves the strand specificity of *intI1* and known *attC*s. Thus, the two tested integrons accept cassettes from both sources, although misoriented relative to other cassettes and the direction of the prevailing integron array promoter, supporting a model of cassette privatization through mutual indecipherability (Fig. 2C).

### Enrichment of counterdefense-related functions in phage-encoded integrons

In light of these data suggesting that phage-encoded integrons are a novel integron subtype specific to phage, next we asked what functions the proteins in these loci have, particularly motivated by recent discoveries of integron-encoded defense systems (*4*–*7*). We annotated the cassette proteins with two complementary phage-specific tools: Phold (*17*), which assigns prokaryotic virus remote homologous group (PHROG) functional categories on the basis of protein structure, and Phynteny-transformer (*18*), which predicts gene function from genomic context. Phold classified about 71% (715/1,002) of cassette proteins as unknown function, compared with 44% (2,827/6,451) of non-cassette proteins from integron-containing genomes. Categories related to virion structure, lysis, nucleotide metabolism, and transcriptional regulation were depleted relative to background genes. In contrast, the “moron, auxiliary metabolic gene and host takeover” category was enriched (Fig. 4A). When we applied Phynteny-transformer to the proteins left unclassified by Phold, the structural, lysis, and transcription regulation categories were again depleted and the auxiliary metabolic gene and host takeover category was again enriched in addition to integration and excision and nucleotide metabolism (Fig. 4B).

**Fig. 4.**
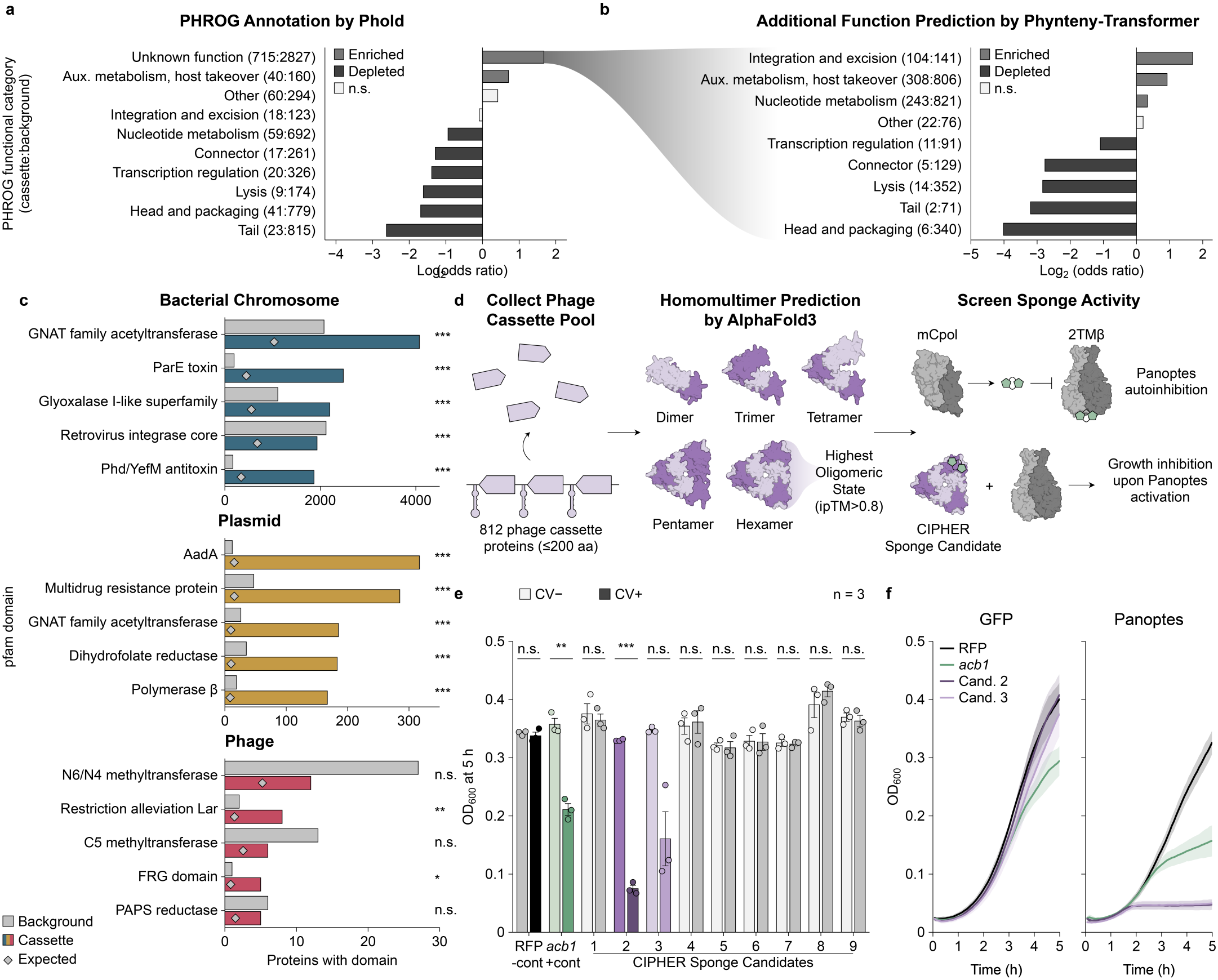
Phage-encoded integrons accumulate counterdefense genes. **(A)** PHROG categories of cassette proteins (Phold): log2 odds ratios versus background proteins (non-cassette proteins of the same genomes); counts (cassette:background) in parentheses. Red, enriched; blue, depleted (FDR < 0.05); gray, not significant (n.s.; Fisher’s exact test, Benjamini-Hochberg correction). **(B)** Phynteny-transformer categories for proteins unclassified by Phold, as in **(A). (C)** Five most frequent Pfam domains in cassette proteins of each dataset: proteins with each domain (cassette, red; background, gray) and number expected without enrichment (diamonds); *, **, ***, FDR < 0.05, 0.01, 0.001. **(D)** Left, AlphaFold3 homomultimer predictions for 812 cassette proteins of fewer than 200 amino acids; nine candidates were synthesized (Fig. S3). Right, Panoptes assay in *E. coli* with an inducible candidate. **(E)** OD_600_ at 5 h with Panoptes and cassette1-9 (Cand. 1-9), Acb1 (positive control), or RFP (negative control), without (CV−) or with (CV+) crystal violet. Mean ± SEM of three biological replicates (circles); \*\**P* < 0.01, \*\*\**P* < 0.001 (paired t test). **(F)** Growth curves (OD_600_) of cells with GFP (no-defense control; left) or Panoptes (right) and RFP, Acb1, cassette2, or cassette3 (Cands.) under crystal violet. Mean ± SD of three biological replicates.

Pfam annotation (*19, 20*) of the cassette proteins from all three datasets gave a more specific view of the gene families enriched relative to background genes (Fig. 4C). The most frequently detected domains in phage cassettes were N6_N4 methyltransferase, restriction alleviation protein Lar, C5 methyltransferase, the FRG domain, and PAPS reductase, of which Lar and FRG were significantly enriched (false discovery rate, FDR < 0.05). Methyltransferases and restriction alleviation proteins are known to function in anti-restriction-modification (*21, 22*). Bacterial and plasmid cassettes, in contrast, were abundant in domains related to antibiotic resistance genes (ARGs), including GNAT family acetyltransferase, glyoxalase, AadA, multidrug resistance protein, and dihydrofolate reductase (DHFR), and in toxin-antitoxin domains such as ParE toxin and Phd/YefM antitoxin, in agreement with previous studies (*11, 23, 24*).

These results suggest that phage-encoded integrons accumulate accessory genes, including counterdefense genes, whereas bacterial and plasmid integrons accumulate antibiotic resistance and toxin-antitoxin genes. To capture a broader set of counterdefense genes, we turned to structure-based modeling, focusing on sponge proteins. These proteins function as counterdefense effectors that neutralize host defense by sequestering immune signaling molecules. Following previous work (*25, 26*), we predicted homomultimers for all cassette proteins smaller than 200 residues using AlphaFold 3 (*27*) to search for sponge-like characteristics, and synthesized nine candidates after visual inspection of their predicted structures (Fig. S3 and Tables S2 and S3).

We assessed these putative sponge proteins with the previously characterized Panoptes system (*28*) (Fig. 4D), which detects the reduction in concentration of nucleotide-derived second messengers by sponge proteins and triggers cell death. In this growth assay, cells co-transformed with the Panoptes system and a putative sponge protein cloned under an inducible promoter are grown in the presence of an inducer. In an endpoint growth screen of all nine candidates co-expressed with Panoptes, induction of cassette2 significantly reduced growth alongside the positive control Acb1 (*P* = 6.4 × 10?^4^ and 0.0013, respectively; paired t-test; Fig. 4E). Cassette3 reduced growth in all three replicates, but to a variable extent, and the difference was not significant (*P* = 0.051); none of the other seven candidates had a significant effect (Fig. 4E). Growth curves with and without Panoptes showed that both cassette2 and cassette3 arrested growth only in its presence of Panoptes (Fig. 4F), providing functional evidence of sponge activity.

## Discussion

In this study, we report that phage-encoded integrons are a distinct subtype of integron with a unique architecture and functionality. Architecturally, their cassettes mostly lie in the reverse orientation compared to non-phage integrons, which privatizes their cassette pool; consistently, most CIPHER genes (96%) lack homologs in non-phage integrons. Evolutionarily, the phage *intI*, like that of class 1 integrons, probably derives from Betaproteobacteria, but appears to have been domesticated independently and possibly much earlier. Functionally, the annotatable fraction of CIPHERs is enriched in counterdefense functions, in contrast to the recently reported defense system biobank of chromosomal and mobile integrons.

Phage-encoded integrons and their CIPHERs shed light on the ecology and evolution of integron cassettes. The strand selectivity of integration provides mutual indecipherability between CIPHERs and cassettes encoded by other integrons (Fig. 2A-C), which insulates productive exchange between the two cassette pools. This separation makes sense given the presumable fitness costs of integrating cassettes of the other pool. More specifically, CIPHERs in bacterial integrons could inhibit or trigger the acquired defense systems of the host, and CI/MI-derived antiphage defense systems in phage-encoded integrons could inhibit the phage life cycle. Although this defense-counterdefense contrast is clear, we cannot conclude that it was the driving force behind this cassette privatization. Other effects, including the toxicity of some phage proteins (or cassettes) within bacterial cells, may cause this difference, meaning the defense-counterdefense contrast itself may be a result and not a cause. Beyond cassette ecology, CIPHERs also provide insight into how genes are captured as cassettes. Most cassettes characterized so far follow the same orientation (forward array), with toxin-antitoxin (TA) cassettes being the exception. The existence of CIPHERs suggests that the capture of a gene by an *attC* site is not biased in orientation, and that the bias instead arises through subsequent selective pressures, when cassettes of different orientations are preferentially fixed in different cassette pools, CI/MI or CIPHERs. Alternatively, CI/MI and phage-encoded integrons might each carry an independent mechanism that generates only cassettes of the corresponding orientation, but there is little data that supports this scenario.

Since phage-encoded integrons are rare in the dataset, the 124 stringently identified elements studied here may not capture their full landscape. Because most cassettes remain unannotated, the proposed role of CIPHERs as a counterdefense biobank rests on the annotated fraction and on two sponge proteins validated with Panoptes. Direct tests of counterdefense during phage infection and large-scale functional assays of CIPHERs will clarify the role of phage-encoded integrons in the phage life cycle and may uncover unknown counterdefense genes and other accessory genes. Lastly, we detected no integrons in isolated phages. Expanding the dataset or the sample collection should allow us to investigate the function and dynamics of phage-encoded integrons in their native context. With these limitations in mind, our findings also provide potential applications for counterdefense gene mining. Whereas antiphage defense systems of bacteria have been characterized extensively and– from restriction enzymes to CRISPR-Cas, widely used as biotechnological tools (*29*–*32*)–counterdefense systems of phages remain far less explored despite their promise for improving phage therapy, microbial community editing, and the treatment of antimicrobial resistance infections. Recent approaches to catalog nature’s vast counterdefense gene reservoir include guilt-by-association (*33*), self-targeting (primarily CRISPR) (*34, 35*), and hypervariable region mining (*36*). CIPHERs add a new genomic signal to flag genes with high counterdefense potential, including among the many cassette proteins that cannot yet be annotated.

More broadly, our findings change the view of the role integrons play in bacteria-MGE conflict and evolution. While integrons have been regarded as the means by which bacteria stockpile defense genes against phages and other mobile genetic elements, our work highlights that the same integron machinery equips phages for the other side of the arms race, with an inverted cassette orientation that keeps the two armories apart.

## Acknowledgements

We thank Sophia Swartz, Pia Andrade, Jack Demaray, Meghna Vergis, Liana Yong, Siqi Yang, Jaymin Patel, and Zach Laturner for critical feedback on this manuscript; Santiago Lopez, Peter Yoon, Kim Seed, and all members of the Cress lab for helpful discussions; Liana Yong, Leo Song, and Santiago Lopez for technical support; and Santiago Lopez for providing plasmids. We gratefully acknowledge the leadership and support of the Berkeley Initiative for Optimized Microbiome Editing (BIOME), particularly Jennifer Doudna, Jill Banfield, Brad Ringeisen, Audrey Glynn, and Rachel Evans. We used Claude Code (Anthropic) and Codex (OpenAI) for coding and writing support.

## Author contributions

Conceptualization: RO, BFC

Experimental Design: RO, BFC

Experimental Work: RO

Computational Analysis: RO

Manuscript Preparation: RO, BFC

Supervision: BFC

## Data, code, and materials availability

Data S1

## Methods

### Datasets and integron detection

We analyzed 166,726 phage genomes with ‘high-quality’ or ‘complete’ annotation from IMG/VR v4 database, downloaded via PhageScope database (*37, 38*), 56,259 complete bacterial genomes from RefSeq (*39*), and 50,553 plasmids from PLSDB, obtained via PlasmidScope database (*40, 41*); the last two served as controls for chromosomal integrons (CI) and mobile integrons (MI), respectively. We ran IntegronFinder v2.0.5 (*42*) with options “--local-max --union-integrases --calin-threshold 1 --promoter-attI --circ”, counting all tyrosine recombinases as integrases to maximize sensitivity, and did not distinguish complete integrons from CALINs (clusters of *attC* sites lacking integrases). We discarded In0 elements, which lack *attC*.

For the bacterial chromosomes, we predicted proviruses with geNomad v1.7.6 (end-to-end, default settings; database v1.7) (*43*), refined their boundaries with CheckV v1.1.1 (end_to_end; database v1.5) (*44*), and discarded any integron element overlapping a provirus by at least 1 bp, irrespective of CheckV quality, which reduced the detected integron count from 16,813 to 16,152. We then retained elements with at least two *attC* sites (CI and MI) or at least three (phage-encoded integrons) (Fig. 1C). We deduplicated phage genomes by viral operational taxonomic unit (vOTU) provided by IMG/VR, and plasmids by MOB-typer (*45*) primary cluster provided by PLSDB, keeping one genome with the most *attC* sites in retained elements from each cluster. This yielded 124 phage-encoded integrons (117 genomes), 534 MI, and 8,972 CI (Fig. 1B), hereafter the filtered datasets.

### Extraction of integron components

We used the *attC* sites called by IntegronFinder without modification. For integrases and cassette proteins, we replaced IntegronFinder’s gene calls with the CDS annotations of the source databases: each ORF called by IntegronFinder was matched to and replaced with a same-strand CDS with at least 80% reciprocal positional coverage from the source database. ORFs without a unique match were discarded. Within each integron element, we paired each *attC* with the nearest cassette protein (edge-to-edge distance); when multiple *attC*s had the same nearest protein, only the closest was paired and the others stayed unpaired. A pair was forward when the cassette CDS lay on the same strand as the top strand (ts) of the *attC*, the strand the *attC* covariance model from IntegronFinder calls, and reverse otherwise (Fig. 2A). To compare the fit of forward and reverse *attC*s to attc_4 (Fig. S2), we used the e-value that IntegronFinder reported for each paired *attC*. IntegronFinder searches each replicon as a whole, so the e-value of an *attC* depends on the length of its replicon, and forward and reverse *attC*s come from replicons of different lengths; we therefore divided each e-value by the length of its replicon in Mb and compared −log10 (e-value per Mb) between forward and reverse *attC*s in each dataset with a two-sided Mann-Whitney U test. Cassette proteins were all proteins of an integron element other than the integrase (n = 1,002 in phage-encoded integrons: 162 forward, 385 reverse, and 455 unpaired). Background proteins were all other proteins from the same genome, for phage and plasmid dataset. For RefSeq, we sampled up to 100 proteins per genome at random (seed 42), excluding proteins overlapping a provirus.

### Homology between cassette pools

We searched the cassette proteins of CI (n = 102,229), MI (n = 2,793), and phage-encoded integrons (n = 1,002) against one another with DIAMOND v2.1.9 (*46*) blastp (--very-sensitive, -k 10). In this search, homologs were defined as a pair of protein with E < 10^−5^, identity ≥ 30%, and reciprocal coverage ≥ 50%.

### Structure prediction and clustering of attC sites

We re-scored the IntegronFinder *attC*s against the covariance model from IntegronFinder (attc_4) with cmsearch (Infernal 1.1.5; -E 10 --max --notrunc) (*47*) to obtain bit scores, which we used to select the phage *attC*s for structural clustering; *attC*s without a hit were not scored. To build a reference set, we scored all 114,655 RefSeq CI *attC*s against the 11 taxon-specific covariance models of previous study (*12*) and kept the 10 highest-scoring *attC*s per model. We excluded three models (Acidobacteria, Deltaproteobacteria, and Methylococcales) for specificity purposes. The remaining eight models gave 80 reference *attC*s, which we grouped into the XL (Xanthomonadales), soil/freshwater (Pseudomonadales, Planctomycetes, Cyanobacteria, and Spirochaetes), and marine (Vibrionales, Alteromonadales, and Oceanospirillales) clades. The 82 phage *attC*s were the reverse-cassette *attC* with the highest attc_4 bit score (≥ 20) from each of 74 vOTUs and all nine *attC*s of IMGVR_UViG_3300010345_000492 (Fig. 2F).

We aligned each *attC* to attc_4 with cmalign (*47*) to locate its conserved boxes and folded the bottom strand (bs) of each of the 162 *attC*s and both strands of the five *attC*s of Fig. 2H as single-stranded DNA with RNAfold (ViennaRNA 2.5.1; dna_mathews2004 parameters; -C --enforceConstraint) (*48*), imposing those of the 14 consensus box base pairs of attc_4 (R″ 1-7/R′ 47-41, L″ 16-18/L′ 30-28, and L″ 20-23/L′ 27-24) that could form in each sequence. At least 10 pairs could be imposed for each of the 162 *attC*s, and all 14 for 40 of them. For the structures of Fig. 2H, a pair was imposed only if both of its boxes had the full width of the model, which left 12, 12, 7, and 6 pairs for ph*attC-1, -3, -5, and -6* (R′ of ph*attC-5* and R″ of ph*attC-6* are one nucleotide short) and 14 for *attC-aadA7*. The extrahelical base (EHB) slot (model column 19) was left unconstrained. We clustered the bs folds with RNAclust v1.3 (LocARNA 2.0.0RC10; UPGMA) (*49*) on the base-pair probability matrices of the constrained folds (ViennaRNA 2.7.2; RNAfold -p -d2 with the same constraints; RNAclust --dpdir) (*48*) (Fig. 2G). We assigned each phage *attC* to the clade of the reference *attC* with the smallest distance in the matrix of pairwise distances underlying the dendrogram.

### Phylogeny of integron integrases

Reference full-length *intI* sequences were retrieved from Supplementary Figure S2 of Cury et al. (*13*). We obtained 175 unique sequences after deduplication, including two XerC/XerD sequences, which were used as an outgroup together with other two XerC/XerD sequences added manually. The 10 phage integrases were those carrying a bona fide *intI* (annotated as ‘intersection_tyr_intI’ by IntegronFinder) in integrons with at least three *attC*s in the filtered phage dataset, irrespective of cassette orientation. Predicted phage hosts were taken from the IMG/VR v5 host annotation. We aligned the 187 sequences with MAFFT v7.505 (L-INS-i) (*50*), trimmed the alignment with trimAl v1.5 (-gt 0.2; 340 columns) (*51*), and inferred a maximum-likelihood tree with IQ-TREE v3.0.1 (-m MFP -B 1000 --alrt 1000; Q.PFAM+I+R7 selected by BIC) (*52*). We rooted the tree at the most recent common ancestor of the four XerC/XerD sequences and considered branches supported when the SH-aLRT was ≥ 95. The tree was visualized in iTOL (*53*) and pruned for display in Fig. 3A.

### Functional annotation of cassette proteins

We annotated phage cassette and background proteins with Phold v1.2.2 (*17*) and, for proteins that Phold left unclassified, with Phynteny-transformer v0.1.3 (*18*), applying no confidence threshold. For Pfam annotation, we searched cassette and background proteins against Pfam 37 (*20*) with hmmscan (HMMER 3.3; -E 0.001) (*19*) and accepted domains with an independent E-value ≤ 0.001 in order of increasing E-value, discarding a hit when at least 20% of its span was already covered by an accepted domain; each protein was counted once per domain. For each domain, we compared the numbers of cassette and background proteins carrying it with a two-sided Fisher’s exact test over all proteins of each group, corrected the *P* values over all domains of the dataset by the Benjamini-Hochberg method, and called a domain enriched when FDR < 0.05 and the odds ratio was above 1. Domains were ranked by the number of cassette proteins, with ties broken by the smaller FDR; this decided the last two places of the phage panel among three domains with five cassette proteins each (FRG, PAPS reductase and a phage integrase domain). The genome in Fig. 2F was annotated with Pharokka v1.9.1 (*54*).

### Structure prediction of cassette proteins

We predicted the structures of all 1,002 phage cassette proteins as monomers and, for the 812 proteins of 200 residues or fewer, as homomultimers (two to six identical chains) with AlphaFold 3 (v3.0.1; seed 42, five models per input, otherwise default settings) (*27*), and used the top-ranked model by ranking score. Following Yirmiya et al. 2025 (*25*) and Tal et al. 2026 (*26*), we assigned each protein the oligomeric state with the most subunits among those with an interface predicted TM-score (ipTM) > 0.8, which applied to 124 of the 812 proteins; we inspected these structures manually and synthesized nine. We searched the monomer models of all 1,002 cassette proteins against the PDB (pdb100, release 2025-01-01) with Foldseek (commit 967df25; easy-search, -- alignment-type 3, -s 9.5, --max-seqs 4000) (*55*). For each protein, the top three hits ranked by the smaller of the TM-scores normalized by the query length and by the target length are shown in Table S3.

### Bacterial strains and media

*E. coli* NEB10β (New England Biolabs) was used as the recipient in the conjugation assay and as the host in the Panoptes assay. *E. coli* DATC (DAP-auxotroph; (*56*)) was used as the donor in the conjugation assay. Cultures were grown in LB at 37 °C with chloramphenicol (34 µg/ml), kanamycin (25 µg/ml) and carbenicillin (100 µg/ml) as required; 300 µM diaminopimelic acid (DAP) was added for all cultures for DATC. All plasmids were verified by whole-plasmid sequencing (Plasmidsaurus). The metadata and sequences of plasmids, oligonucleotides and gBlocks used are listed in Tables S4 and S5.

### Suicide conjugation assay

To measure cassette insertion, we used a suicide conjugation assay (*16, 57*) (Fig. 3B). The *attC* sites tested (*attC-aadA7* and ph*attC-1, -3, -5 and -6*, which are *attC* 1, 3, 5 and 6 of the array in Fig. 2F; Table S1) were synthesized as oligonucleotide pairs with 3-nt overhangs, annealed, and cloned in both orientations by LguI Golden Gate assembly into a PCR-amplified backbone of pRO0054, a plasmid with an R6K origin, an oriT and a kanamycin resistance gene. Because the strand transferred from the oriT of pRO0054 is fixed, the two orientations of an insert deliver the top strand (ts) or the bottom strand (bs) of the attC to the recipient as single-stranded DNA (donor plasmids for each strand in Table S1). The plasmids were electroporated into *E. coli* DATC, which expresses the π protein required for R6K replication, and these strains served as donors. Recipients were NEB10β, co-transformed with an *attI* plasmid and an integrase plasmid by heat shock (as described for the Panoptes assay below). The *attI* plasmid was made from pBFC0011 (p15A origin, CmR, GFP) by replacing GFP with an *attI* sequence, synthesized as a gBlock (IDT), by BsmBI Golden Gate assembly. For class 1 integrons we used the 90-bp *attI1* sequence shown in Fig. 1b of (*58*), and, as the putative *attI* of the phage integron, the 120-bp intergenic region between *intI* and the first cassette gene of the element in Fig. 2F (positions 29,393-29,512 of IMGVR_UViG_3300010345_000492). The integrase plasmid was made from pRO0005 (SC101 origin, AmpR, mKate2 under the pJEX promoter) by replacing mKate2 with an *intI* gene by PaqCI Golden Gate assembly, which places *intI* under the crystal violet-inducible pJEX promoter. IntI genes were synthesized as a gBlock (IDT). The phage *intI* (positions 28,367-29,392) carries three silent substitutions that remove internal PaqCI sites; *intI1* was taken from NZ_JBSPSX010000414.1, where it encodes WP_000845048.1 (*59*). pRO0005 itself served as the no-integrase control.

Donor and recipient overnight cultures were grown at 37 °C with the appropriate antibiotics (donors: kanamycin and 300 µM DAP; recipients: chloramphenicol and carbenicillin). Cells were collected (4,000 g, 10 min), resuspended in 1 ml LB, washed three times in LB, and resuspended in 1 ml LB for measuring OD_600_. After a further centrifugation, cells were resuspended in LB containing 300 µM DAP and 250 nM crystal violet at an OD_600_ of 10 (about 600 µl). Donor and recipient suspensions (50 µl each) were mixed in a 96-well plate, and 10 µl of each mixture was spotted with a multichannel pipette onto LB agar containing 300 µM DAP and 250 nM crystal violet. After 4 h of conjugation at 37 °C, each spot was scraped into 100 µl LB in a 96-well plate, serially diluted 10-fold, and 5 µl of each dilution was spotted onto LB agar containing chloramphenicol (to count recipients) or kanamycin (to count transconjugants), without DAP. Colonies were counted after about 20 h at 37 °C. The insertion efficiency was calculated as KanR CFU divided by CmR CFU, using the spot of the dilution at which colonies could be counted; a sample without KanR colonies was scored as zero (limit of detection: one colony in the undiluted KanR spot). Data are the mean ± s.e.m. of three biological replicates, each started from independent overnight cultures on a different day. Representative transconjugant colonies selected for plasmid recombination were sequenced by Plasmidsaurus whole-plasmid sequencing to confirm the intended recombination and to determine the insertion orientation.

### Panoptes growth assay

The Panoptes plasmid pSCL1237 (p15A origin, CmR) carries the Panoptes locus under its native promoter, with an upstream pTet promoter that was left uninduced throughout. pBFC0011 (p15A origin, CmR, GFP) was used as the negative control. Candidate genes were synthesized as gBlocks Gene Fragments (IDT) and cloned by Golden Gate assembly with PaqCI (NEB) into pBFC1299 (SC101 origin, KanR), replacing mKate2 downstream of the crystal violet-inducible pJEX promoter (pRO0031-pRO0039; proteins listed in Table S2). pBFC1299 retaining mKate2 (RFP) served as the cassette-negative control. As a positive control, pRO0029, which carries Acb1 of *Erwinia* phage FBB1 (UniProt A0A868BQY3) in the same backbone was used.

The Panoptes/GFP plasmid and a candidate/mKate2 plasmid were co-transformed into chemically competent *E. coli* NEB10β cells (New England Biolabs): 10 ng of each plasmid was added to one tube of cells, kept on ice for 20 min, heat-shocked at 42 °C for 45 s, returned to ice for 2 min, and recovered in 450 µl SOC at 37 °C for 1 h. Cells were plated on LB agar with chloramphenicol and kanamycin, and single colonies were picked and stored as glycerol stocks. Overnight cultures were started from the stocks in LB containing both antibiotics, and each biological replicate was started from an independent overnight culture. 8 × 10^6^ CFU were seeded per well in 200 µl of LB containing both antibiotics and crystal violet (0 or 100 nM; 70 nM for Acb1 (maximum induction level before intrinsic Acb1 toxicity)) in a 96-well plate. OD_600_ was measured every 5 min for 16 h at 30 °C with continuous orbital shaking (282 cpm, 3 mm) in a BioTek Synergy H1 plate reader, and the signal of wells containing medium only was subtracted as the blank. For Fig. 4E, the OD_600_ at the read closest to 5 h of the wells with and without crystal violet was compared for each strain across the three biological replicates with a two-sided paired t-test, without correction for multiple comparisons.

## Supplementary Materials

Tables S1 to S5

## Supplementary Figures

**Fig. S1.**
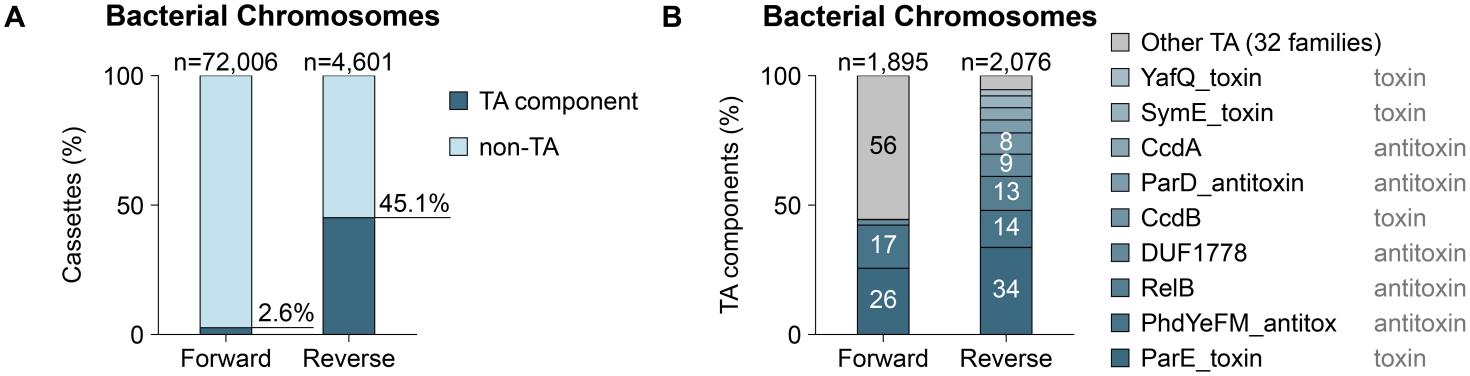
Toxin-antitoxin components among the forward and reverse cassettes of chromosomal integrons. **(A)** Percentages of the cassette proteins paired with an *attC* in bacterial chromosomes (n = 72,006 forward and 4,601 reverse cassettes) that are toxin-antitoxin (TA) components, that is, that carry a Pfam domain of a TA family (Pfam 37 families whose description contains toxin-antitoxin, antitoxin, antidote, addiction module, or post-segregation, plus CcdB, RES, HicB, MqsR, the three RnlA toxin domains, and DUF1778). Pfam families that also occur in many non-TA proteins (for example GNAT acetyltransferase, PIN, Fic/Doc, and HTH_3) were not counted, so the percentages are lower bounds. **(B)** Composition of the TA components in **(A)** (n = 1,895 forward and 2,076 reverse) by Pfam family; nine families are shown and the other 32 are pooled.

**Fig. S2.**
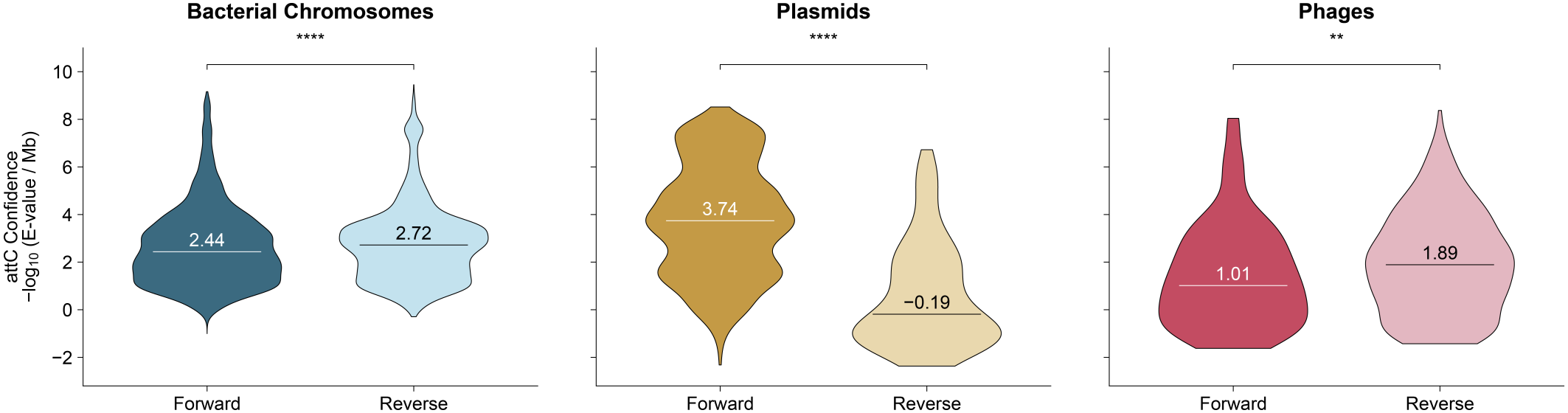
Fit of the *attC*s of forward and reverse cassettes to the *attC* covariance model. Distribution of the fit of *attC*s to attc_4, shown as −log10 (E-value per Mb of the searched replicon; larger values indicate a better fit), for *attC*s paired with forward or reverse cassettes in bacterial chromosomes (n = 72,006 forward and 4,601 reverse), plasmids (n = 1,700 and 112), and phages (n = 162 and 385). Lines and numbers indicate medians. **, *P* < 0.01; ****, *P* < 0.0001 (two-sided Mann-Whitney U test).

**Fig. S3.**
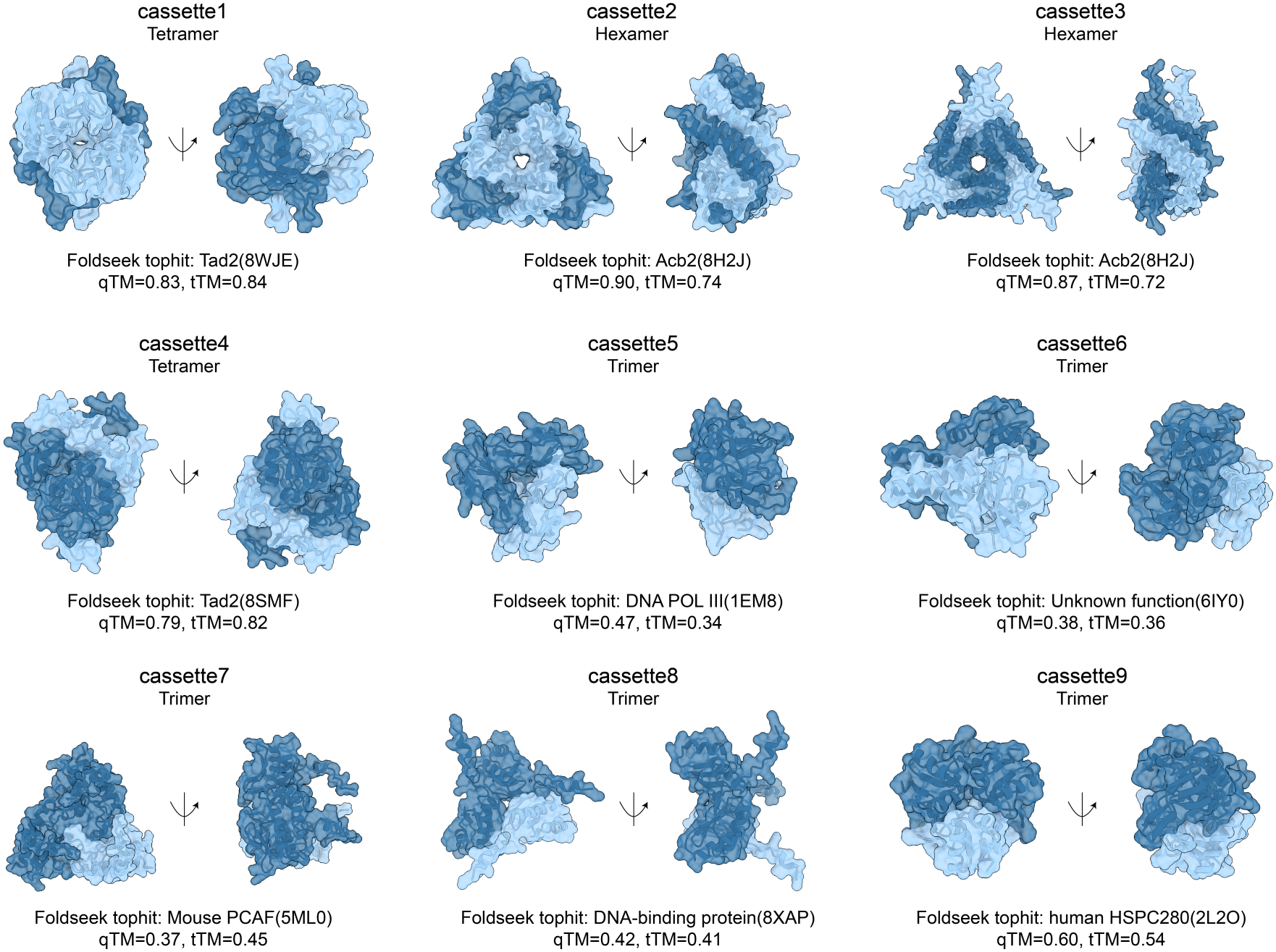
AlphaFold3 models of the nine candidate sponge proteins. Each candidate (cassette1-9) is shown in its oligomeric state (the largest oligomer with ipTM > 0.8). Below each model, the top hit of a Foldseek search of the monomer model against the PDB (pdb100) is given with its TM-scores normalized by the query (qTM) and by the target (tTM). For each of the 1,002 cassette proteins, the top three hits ranked by the smaller of qTM and tTM are listed in Table S3.

